# An AmotL2–Yap1 Module Integrates Flow and Junctional Mechanics to Specify Vascular Pruning Hotspots

**DOI:** 10.1101/2025.11.19.689183

**Authors:** Maria P. Kotini, Ludovico Maggi, Etienne Schmelzer, Hiroyuki Nakajima, Heinz-Georg Belting, Markus Affolter

## Abstract

Vascular networks constantly remodel to optimise blood flow, yet how endothelial cells (ECs) integrate haemodynamic cues with junctional mechanics to execute precise branch pruning remains unclear. Using live imaging and quantitative topology analysis in the zebrafish sub-intestinal vein plexus, we define the cellular events underlying flow-guided regression. Pruning occurs in geometrically disadvantaged, low-flow branches and is executed through transient polarity fluctuations, junctional remodelling and directed EC migration. We identify AmotL2 as an essential regulator of endothelial mechanical competence. AmotL2 maintains junctional architecture, actomyosin tension and appropriate cell packing, enabling ECs to polarise and rearrange in response to local flow differences. Loss of AmotL2 disrupts tension homeostasis, increases tissue compaction and prevents branch regression despite normal perfusion. In contrast, Yap1 plays an opposing, stabilising role, and its loss results in excessive pruning and network simplification. Together, AmotL2 and Yap1 constitute a mechanosensitive balancing module that interprets flow magnitude and vascular geometry to determine branch stability. This framework describes how mechanical forces sculpt vascular architecture during development and homeostasis.

## Introduction

Vascular networks must be continuously refined to meet the oxygen and metabolic demands of developing tissues. Angiogenesis establishes the initial blood vessel patterns (Gebala et al., 2016; Hogan and Schulte-Merker, 2017; Phng and Belting, 2021), and subsequently vascular remodelling determines whether these nascent networks become functional through network optimisation. During this optimisation phase, high-flow branches are stabilised, while redundant or inefficient segments are eliminated (Furtado and Eichmann, 2024; Korn and Augustin, 2015). Remodelling and network refinement occurs across organs, including the retina (Franco et al., 2015), the skin (Kam et al., 2023) and the brain (Chen et al., 2012; Kochhan et al., 2013), where coordinated endothelial rearrangements and branch regressions sculpt mature blood vessel patterns. Blood flow is a central regulator of these behaviours, where local haemodynamic differences determine whether endothelial cells (ECs) maintain junctional stability, migrate, or exit a branch entirely via pruning (Chen et al., 2012; Franco et al., 2015; Lenard et al., 2015; Lucitti et al., 2007; Phng and Belting, 2021). In several microvascular beds, including the zebrafish sub-intestinal vein plexus (Lenard et al., 2015) and the mammalian retina (Franco et al., 2015), blood vessel regression is mediated not by apoptosis but by directed migration of endothelial cells (ECs) from poorly perfused branches into adjacent, highly-perfused ones. These observations reveal a conserved principle, where flow heterogeneity establishes local differences in endothelial plasticity, enabling networks to remove superfluous connections while reinforcing functional routes (Korn and Augustin, 2015). This dependence on flow-mediated cell rearrangements leads to the question of how ECs sense and interpret such spatially restricted mechanical cues to achieve precise remodelling decisions.

Flow sensing and mechanotransduction in ECs rely on specialised molecular pathways that translate haemodynamic inputs into coordinated cellular outputs. Both direct sensors and downstream mediators of mechanotransduction are crucial for the adaptation of endothelial cells to mechanical cues. Within this framework, Piezo1-mediated calcium influx enables rapid flow detection and contributes to vascular development and stabilisation (Li et al., 2014; Liu et al., 2020). Shear stress induces EC alignment, polarity and directional migration through a shear stress dependent modulation of adherens junctions that promotes network stabilisation (Conway et al., 2013; Malek and Izumo, 1996; Tzima et al., 2005; Vion et al., 2021). Longer-term mechanotransduction responses involve transcriptional regulators such as YAP/TAZ, which translocate to the nucleus upon flow initiation to coordinate gene-expression programmes supporting polarity, cytoskeletal organisation and vessel stability (Duchemin et al., 2019; Nakajima et al., 2017). Mechanical load reorganises cortical actin and junctional scaffolds, enabling cells to polarise, migrate and redistribute forces across the network (Kondrychyn et al., 2020). Junctional organisation is highly sensitive to changes in flow magnitude and direction, with altered haemodynamics rapidly triggering junctional turnover and collective rearrangements (Lenard et al., 2015; Tzima et al., 2005). Within this framework, AmotL2 emerges as a key junctional scaffolding role that links cadherin-based adhesions to actomyosin networks (Hultin et al., 2014; Hultin et al., 2017) and couples flow modulation to YAP localisation (Nakajima et al., 2017). Under low or absent flow, YAP interacts with AmotL2 at junctions, while with increasing shear, this association loosens, enabling YAP nuclear entry (Nakajima and Mochizuki, 2017; Nakajima et al., 2017). Although components of these pathways have been linked to flow sensing or mechanotransduction, how they integrate to execute the structural transitions that drive branch pruning remains unresolved. In particular, the mechanisms that couple flow sensing to the junctional remodelling and local cell rearrangements required for regression are still poorly understood.

The zebrafish sub-intestinal vein plexus (SIVP) is an ideal system to dissect these interactions and investigate the principles of vascular remodelling. Unlike more stereotyped vascular beds, the SIVP develops into a highly variable plexus without obvious pre-imposed patterning templates, relying instead on emergent behaviours of migrating and proliferating ECs (Goi and Childs, 2015). Early plexus formation is orchestrated by BMP-dependent collective migration from the posterior cardinal vein and by VEGF-dependent directional shifts that generate venous and arterial territories (Hen et al., 2015). As the plexus expands over the yolk, sprouting, proliferation and dynamic filopodia- and lamellipodia-based exploration promote broad vascular extension (Goi and Childs, 2015). Remodelling begins shortly after flow initiation, marked by flow-dependent pruning events that progressively refine the network (Lenard et al., 2015). Although signals controlling initial plexus formation have been described, including Vegf, BMP, Plexin/Neuropilin pathways and soluble decoy receptors (Goi and Childs, 2015; Hen et al., 2015), the molecular machinery controlling later remodelling stages remains largely unexplored. In particular, how ECs integrate haemodynamic forces with mechanosensitive regulators to choreograph the local cell rearrangements that control branch pruning and unstable connections has not been elucidated.

Here, we combine quantitative topology analysis, high-resolution live imaging and genetic perturbations to define the cellular and molecular logic of flow-dependent remodelling in the SIVP. We identify the structural features that predispose specific regions to undergo pruning, characterise the local endothelial rearrangements that resolve unstable connections and uncover the mechanosensitive role of the AmotL2–Yap axis in balancing stability and remodelling. Together, these findings reveal how ECs integrate network topology, haemodynamic cues and junctional dynamics to coordinate the removal of redundant branches and sculpt functional vascular architecture.

## Results

### Flow-Dependent Morphogenetic Phases Define Intestinal Vascular Remodelling

Vascular remodelling within the sub-intestinal vein plexus (SIVP) has previously been shown to depend on blood flow (Lenard et al., 2015). While previous studies have linked pruning to haemodynamic forces (Chen et al., 2012; Franco et al., 2015; Lenard et al., 2015; Phng and Belting, 2021), how flow interacts with vascular geometry to coordinate distinct morphogenetic events remains unclear. To dissect the cellular and haemodynamic basis of intestinal vascular remodelling, we systematically characterised the temporal sequence of morphogenetic processes shaping SIVP development. Time-lapse imaging of *Tg(fli1a:EGFP)* embryos revealed that SIVP network formation proceeded through distinct, but overlapping phases of sprouting (36–50 hpf), anastomosis (38–60 hpf) and remodelling (48–80 hpf) (Fig. 1A-B). Quantitative analysis across developmental time confirmed that remodelling was dominated by two counteracting processes, pruning of low-flow branches and collateral fusions between perfused segments, together refining the vascular network and driving tissue extension (Fig. 1C). Dual imaging of endothelial cells and erythrocyte flow using *Tg(fli1a:EGFP; gata1a:dsRed)* demonstrated that pruning events consistently occurred in branches experiencing reduced or disturbed flow, whereas vessel segments maintaining or increasing flow underwent fusion (Fig. 1D). Geometric analysis further indicated that L-shaped vessel configurations correlated with pruning, while O-shaped geometries were associated with fusion events (Fig. 1E-F), suggesting that vessel topology is a determining factor of remodelling outcomes. Thus, blood flow acts upon the intrinsic network geometry regulating whether branches are stabilised or eliminated. Pharmacological flow reduction using tricaine abolished pruning, while preserving L-type vessels, confirming the requirement for haemodynamic input in vascular refinement (Fig.1G). Together, these data establish the temporal sequence and flow dependence of morphogenetic events underlying intestinal vascular remodelling.

**Figure 1.**
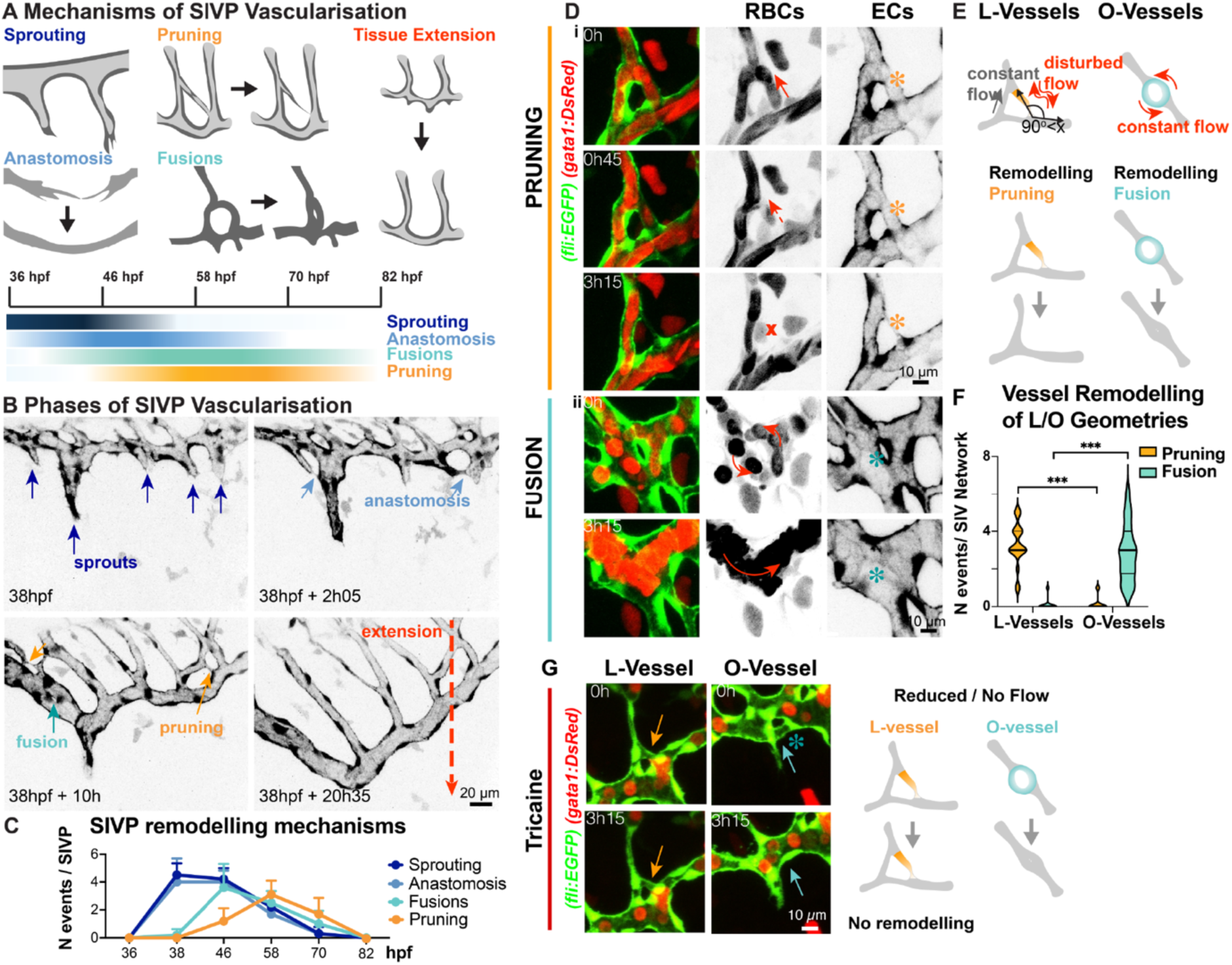
Flow-dependent pruning and fusion shape intestinal vascular remodelling. (A) Schematic illustrating phases of sub-intestinal vein plexus (SIVP) vascularisation, including sprouting (36–50 hpf), anastomosis (38–60 hpf) and remodelling (38–80 hpf), which collectively coordinate tissue extension. Remodelling involves pruning of low-flow branches (42–80 hpf) and collateral fusions (38–76 hpf) that reinforce perfused segments. (B) Time-lapse imaging of SIVP formation in *Tg(fli1a:EGFP)* embryos from 38 hpf. (C) Quantification of morphogenetic events over time (line plot showing mean ± SEM, n = 15 SIVPs from 15 embryos, N= 3 independent experiments). (D) Blood flow dynamics during pruning and fusion in *Tg(fli1a:EGFP;gata1a:dsRed)* embryos at 58 hpf. (i) Vessel branch exhibiting reduced flow prior to pruning. (ii) Vessel segments maintaining constant flow that fuse, leading to increased flow. Red arrows indicate flow direction, orange and cyan asterisks mark pruning and fusion events, respectively. (E) Association between vessel geometry with remodelling outcomes: L-type geometries correlate with pruning, while O-type geometries correspond to fusion. (F) Quantification of L- and O-type geometries and corresponding pruning or fusion frequencies (violin plots displaying data distribution with median and quartiles, n = 20 SIVPs, N = 3 independent experiments; Mann-Whitney U test, ***p < 0.001). (G) Under tricaine-induced blood flow reduction, L-type and O-type vessel branches show absence of pruning, but persistence of fusion in *Tg(fli1a:EGFP;gata1a:dsRed)* embryos at 58 hpf. Orange and cyan arrows indicate L- and O-type vessel geometries, respectively.

### Amotl2 Controls Flow-Dependent Pruning in the Developing Intestine

Given that vascular remodelling relies on flow-dependent branch regression, we next sought to identify molecular regulators linking haemodynamic cues to pruning behaviour. AmotL2 has been implicated in mechanotransduction and junctional regulation (Huang et al., 2007; Hultin et al., 2014; Hultin et al., 2017; Nakajima et al., 2017), prompting us to examine its role in intestinal vascular remodelling. By 82 hpf, at the end of the remodelling phase, control embryos displayed a simplified, hierarchical sub-intestinal vein plexus, whereas *amotL2a/b* double mutants retained a highly complex, unresolved network, indicating defective remodelling (Fig. 2A). Time-lapse imaging revealed that control L-type branches regressed over time, simplifying the plexus, while *amotl2a/b* mutants failed to regress, maintaining aberrant vascular complexity (Fig. 2B). Quantification confirmed a significant reduction in pruning frequency and tissue extension in *amotl2a/b* embryos, while fusion frequency remained largely unaffected (Fig. 2C). Erythrocyte flow analysis revealed normal perfusion dynamics in *amotL2a/b* mutants, excluding impaired blood flow as the primary cause of defective remodelling (Fig. 2D). These findings indicate that AmotL2 specifically regulates pruning downstream of haemodynamic cues rather than flow generation itself.

**Figure 2.**
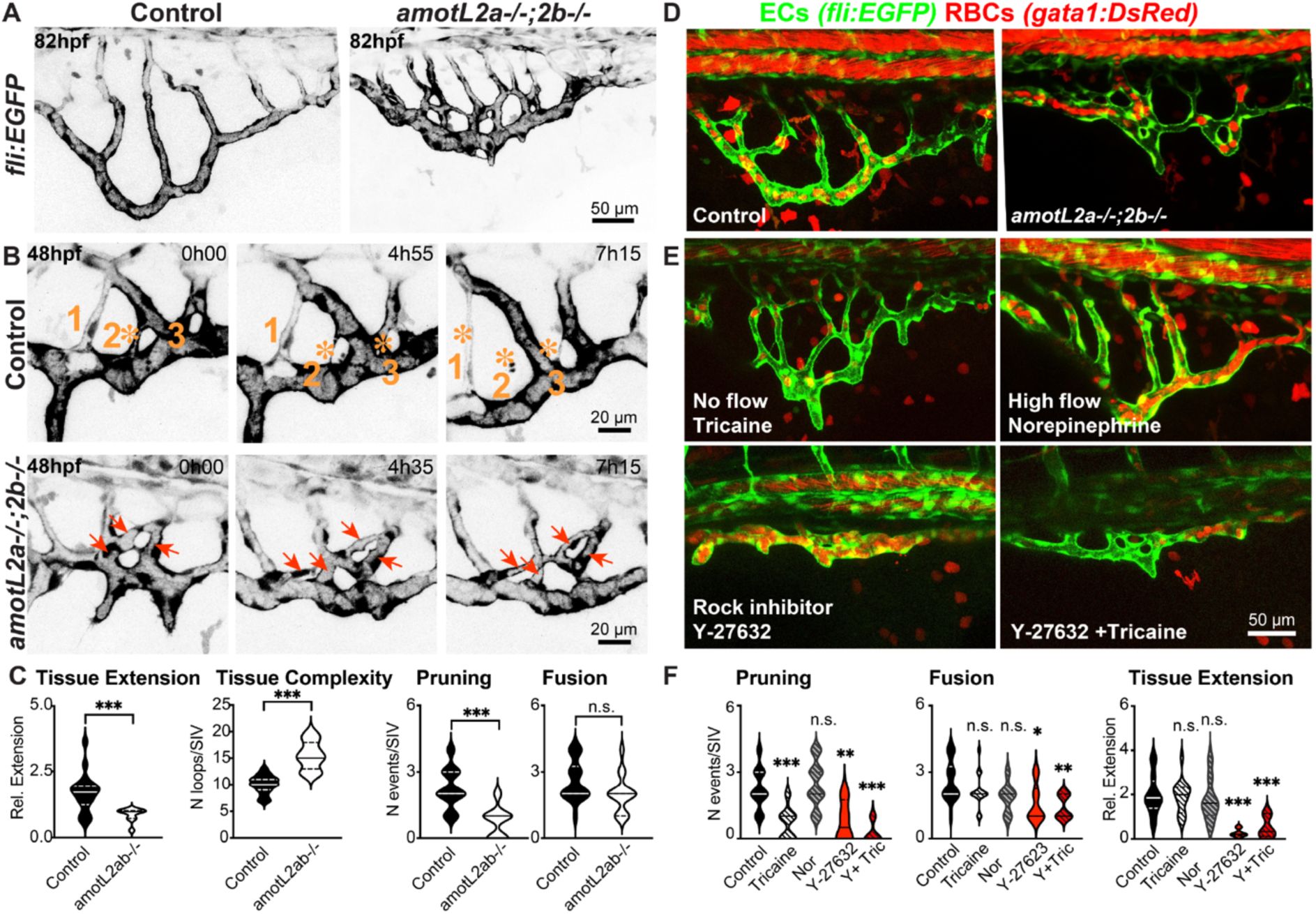
Loss of AmotL2 disrupts pruning and tissue extension, while maintaining vascular complexity. (A) Confocal images of 82 hpf *Tg(fli1:EGFP)* embryos showing simplified vascular networks in controls and increased network complexity in *amotL2a/b* double mutants after completion of remodelling. (B) Representative time-lapse sequences of pruning events at L-type geometries in control embryos and persistent, unresolved branches in *amotL2a/b* mutants starting at 48 hpf. Numbers denote individual branches, orange asterisks indicate pruning events; red arrows mark L-type branches that fail to prune over time. (C) Quantification of tissue expansion, vascular complexity, pruning, and fusion events at 82 hpf (violin plot shoing data distribution, median and quartile range, n = 18 SIVP, 18 embryos, N = 3 independent experiments; Mann–Whitney U test, ***p < 0.001). (D) Confocal images from 82 hpf demonstrates normal blood flow in both control and *amotL2a/b* mutant embryos *Tg(fli1:EGFP; gata1:DsRed)*. (E) Effects of pharmacological perturbations on SIVP architecture in *Tg(fli1:EGFP; gata1:DsRed)* embryos at 70 hpf. Embryos were treated at the peak of SIVP remodelling at 58hpf and during imaging with tricaine (reduced flow), norepinephrine (increased flow), Y-27632 (ROCK inhibitor, reduced actomyosin contractility) or tricaine + Y-27632. Inhibition of remodelling was observed under tricaine and rock inhibition, while tissue extension was reduced only with ROCK inhibition. (F) Quantification of tissue extension, pruning and fusion events following pharmacological treatments (violin plots displaying data distribution with median and quartiles, n = 15 SIVP, N = 6 independent experiments; Kruskal–Wallis test, *p < 0.05, **p < 0.01, ***p < 0.001).

To further probe this relationship, we applied pharmacological treatments modulating flow and contractility. Pruning was strongly reduced by tricaine-mediated flow inhibition and by the ROCK inhibitor Y-27632, which suppresses actomyosin contractility (Fig. 2E). Combined tricaine and Y-27632 treatment nearly abolished pruning, while norepinephrine-induced flow enhancement had minimal effects on pruning or fusion. Tissue extension was similarly decreased under Y-27632 and combined tricaine and Y-27632 treatment, but not under tricaine application alone (Fig. 2F). These findings indicate that AmotL2 specifically mediates pruning downstream of flow-dependent and contractility-dependent cues, coordinating both endothelial branch regression and plexus extension. Thus, coordination between branch elimination and extension is essential for achieving refinement and optimisation of vascular networks.

### Amotl2 Coordinates Junctional Organisation and Cell Density to Permit Flow-Dependent Remodelling

To determine how AmotL2 regulates endothelial behaviour during remodelling, we examined junctional architecture, cell packing and branch morphology. Immunostaining for the tight junction marker ZO-1 revealed that *amotL2a/b* mutants exhibited irregular, highly branched junctional patterns, in contrast to the linear and continuous junctions observed in controls (Fig. 3A), indicating defective cell-cell organisation. To quantitatively assess tissue packing, we measured local endothelial cell spacing in *Tg(fli1a:NLS-mCherry)* embryos using nearest-neighbour relationships derived from Delaunay triangulation. The resulting triangular areas, which inversely reflect local cell density, were significantly reduced in *amotL2* mutants (Fig. 3B–C), indicating increased endothelial compaction. While control SIVPs begin more compact and then progressively increase cell spacing while extending, *amotL2* mutants remain compact throughout development (Fig. 3D). Pharmacological inhibition of ROCK in *Tg(fli1a:pecam1-EGFP; fli1a:NLS-mCherry)* embryos phenocopied the compact morphology seen in *amotL2* mutants, while tricaine treatment did not alter packing (Fig. 3E-F). These results indicate that AmotL2 sustains appropriate endothelial spacing and promotes vascular elongation by maintaining actomyosin-dependent contractility mechanical tension.

**Figure 3.**
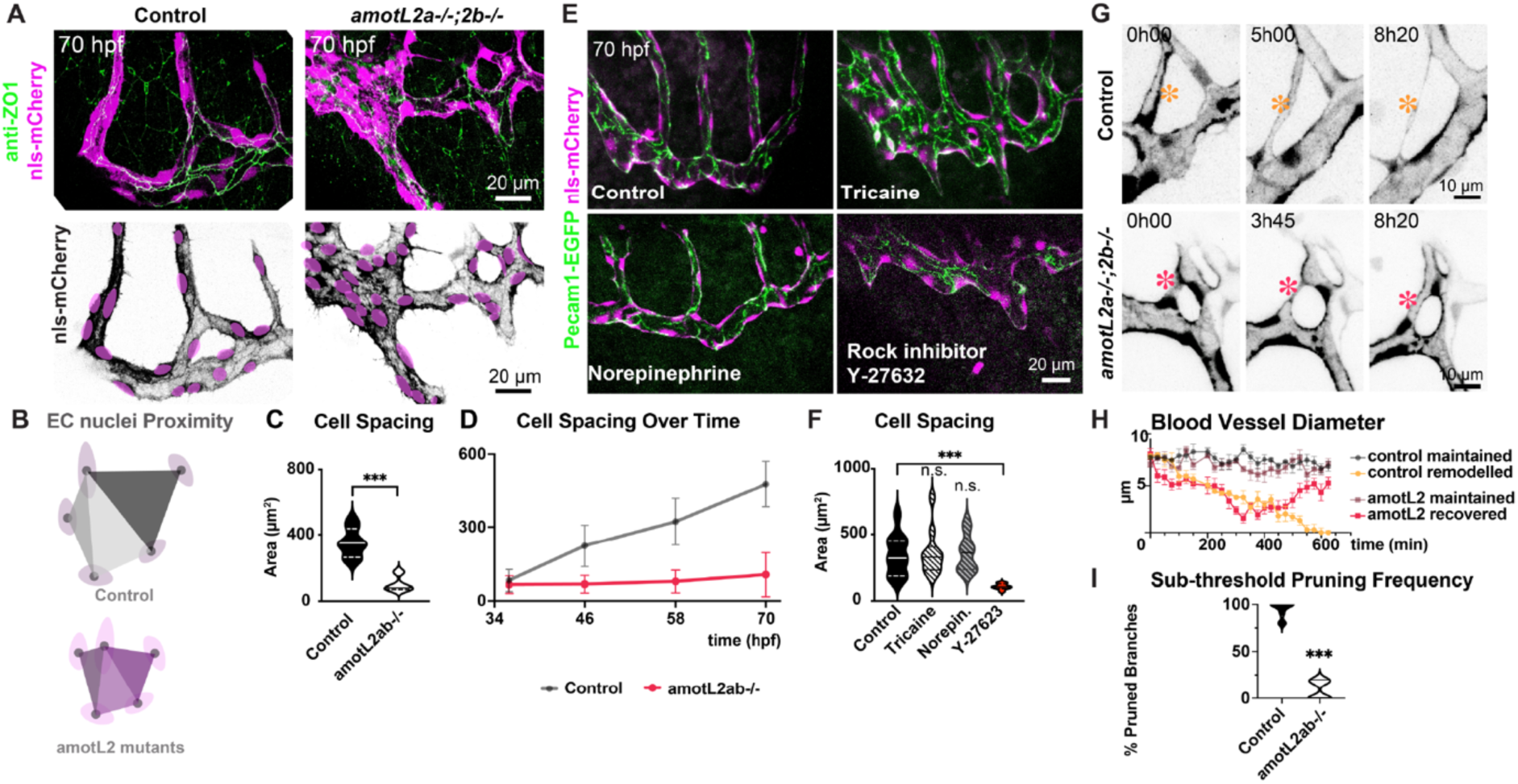
AmotL2 controls endothelial compaction and diameter dynamics during remodelling. (A) Immunostaining of ZO-1 in *Tg(fli1a:NLS-mCherry)^ubs10^* embryos at 70 hpf. *amotL2a/b* mutants display more complex and irregular ZO-1 junctional arrangements within vascular branches compared with controls. Nuclei are pseudo-coloured magenta (NLS-mcherry channel). (B) Schematic illustrating cell proximity metric derived from Delaunay triangulation. Nuclear centroids are connected to their nearest neighbours to form triangular units. The area of each triangle inversely reflects local cell density, with smaller areas indicating increased endothelial compaction. (C) Quantification of endothelial cell proximity at 70 hpf. *amotL2a/b* mutants exhibit significantly reduced cell spacing compared with controls, indicating higher local cell density & tissue compaction (n = 10 SIVPs from 10 embryos, N = 3 independent experiments, Mann–Whitney U test, ***p < 0.001) (D) Analysis of cell density over time demonstrates that control SIVPs progressively decrease in cell density, which correlates with increased tissue expansion, whereas *amotl2a/b* mutants remain compact with reduced extension (n = 8 embryos, N = 3 independent experiments, Mann–Whitney U test, ***p < 0.001). (E) Pharmacological perturbations in *Tg(fli1a: pecam1a-EGFP; NLS-mCherry)* embryos at 70 hpf reveal increased cell compaction following Y-27632 or combined Y-27632 + tricaine treatment, but not with tricaine or norepinephrine alone. (F) Quantification of cell spacing in pharmacologically treated embryos at 70 hpf (n = 10 SIVPs from 10 embryos, N = 6 independent experiments, Mann–Whitney U test, ***p < 0.001). (G) Diameter dynamics in *Tg(fli1a:EGFP)* embryos at 58 hpf show that control vessel branches progressively constrict until reaching a pruning threshold, after which regression ensues. In contrast, *amotl2a/b* branches constrict beyond this threshold, but fail to regress and subsequently recover. Orange asterisk indicates a pruning event in control, red asterisk marks a recovering branch in *amotL2* mutants. (H) Quantification of diameter changes in four groups (control maintained, control pruned, *amotL2* mutant maintained and *amotL2* mutant recovered) demonstrates that *amotL2* mutants exhibit transient constriction without regression (line plot showing mean ± SEM, n = 10 vessels from 10 embryos, N = 3 independent experiments). (I) Proportion of branches that crossed the pruning diameter threshold and subsequently regressed (violin plots displaying data distribution with median and quartiles, n = 10, N = 3 independent experiments, Mann–Whitney U test, ***p < 0.001). The threshold is defined as mean diameter immediately prior to regression in control-pruned branches (4 µm).

Consistent with this, dynamic diameter measurements showed that, control pruning branches gradually constricted towards a reproducible pruning threshold (∼4 µm), after which regression occurred. In contrast, *amotL2* mutant branches frequently constricted below the pruning threshold, but failed to regress and subsequently recovered (Fig. 3G-I). In control embryos, this threshold corresponds to the mechanically weakened, low-flow state that precedes branch removal, consistent with previous observations in brain vasculature (Chen et al., 2012). Together, these findings reveal that AmotL2 couples junctional remodelling and actomyosin-mediated tension to maintain endothelial spacing and enable flow-dependent diameter control, therby ensuring successful branch regression. This findings support that increased tissue compaction in mutants arises primarily from impaired contractility, while impaired diameter dynamics reflect disrupted integration of flow and mechanical tension.

### AmotL2 Controls Endothelial Polarity and Cell Rearrangements during Flow-driven Remodelling

To determine how the absence of AmotL2 causes failure in branch regression, we analysed cell polarity and cell rearrangements within the SIVP. Using *Tg(kdrl:NLS-EGFP; fli1a:B4GALT1-mCherry)* embryos, Golgi orientation relative to flow was used as a proxy for endothelial polarity, as established previously (Kwon et al., 2016). In control embryos, endothelial cells polarised against flow in stable branches and exhibited dynamic polarity fluctuations in pruning branches, reflecting continuous adaptation to local flow changes (Fig. 4A-C). In contrast, *amotL2a/b* mutants displayed restricted polarity dynamics that failed to align with flow, resembling tricaine-treated embryos (Fig. 4A-C). The reduced polarity variance (Fig. 4D) indicates impaired responsiveness to haemodynamic cues, consistent with defects in flow sensing and reduced pruning.

**Figure 4.**
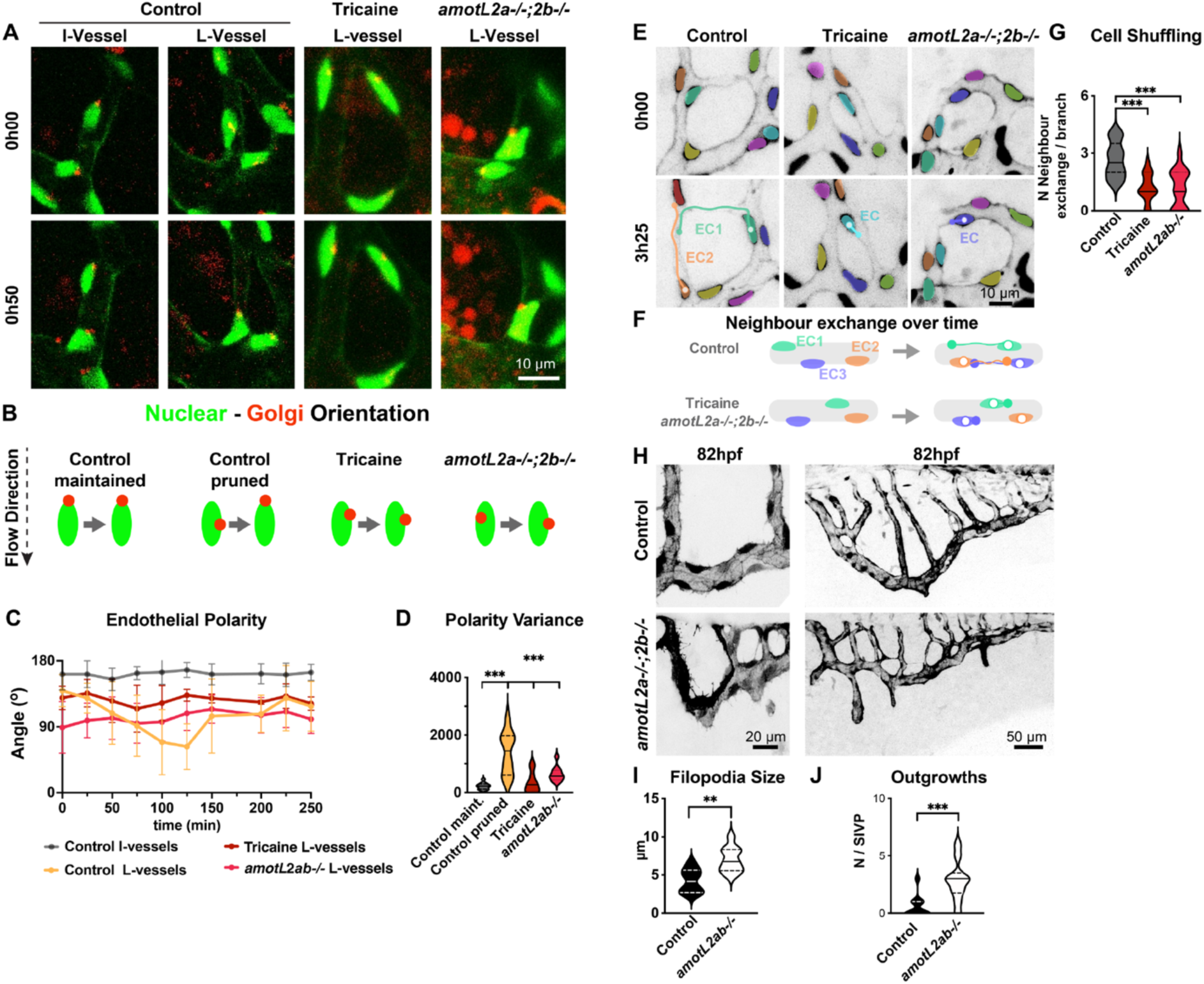
AmotL2 regulates endothelial polarity and cell shuffling during vascular remodelling. (A) Time-lapse snapshots of *Tg(kdrl:NLS-EGFP; fli1a: B4GALT1-mCherry)* embryos at 58 hpf used to assess endothelial cell polarity. (B) Golgi position (red) in relation to nucleus (green) seerves as a proxy for cell polarity relative to flow (black arrow). (C) Quantification of the Golgi-nuclear angle shows that control endothelial cells polarise against flow direction (≈180°) in stable branches (control I-vessels). Polarity fluctuates dynamically in pruning branches (control L-vessels), whereas tricaine-treated embryos display more reduced and misaligned polarity (≈125°). In amotL2a/b mutants, polarity fluctuates within a restricted range (≈95°), indicating defective flow-dependent polarisation (line plot showing mean ± SD, n = 10 embryos, 5 cells per embryo, N = 3 independent experiments). (D) Polarity variance demonstrates greater fluctuation in control-pruned branches compared with control maintained, tricaine-treated or *amotL2a/b* mutant branches (violin plots displaying data distribution with median and quartiles, n = 10 embryos, 5 cells per embryo were analysed, N = 3 independent experiments, Mann–Whitney U test, ***p < 0.001, all comparisons vs. control-pruned branches). (E) Time-lapse snapshots of *Tg(kdrl:NLS-EGFP)* embryos at 58 hpf show reduced endothelial cell shuffling in *amotL2a/b* mutants compared with controls. Pseudocolor-coded cells highlight intercalations in controls and reduced cell shufflling in tricaine-treated and mutant embryos. (F) Neighbour exhange per branch was used as a measure to quantify cell shuffling. Cyan, ornage and purple mark different endothelial cells (ECs). (G) Quantification of neighbour exchange over time shows reduced cell shuffling rates in reduced in *amotL2a/b* mutants and tricaine-treated embryos (violin plots displaying data distribution with median and quartiles, n = 15 embryos, 10 branches per embryo, N = 3 independent experiments, Mann–Whitney U test, ***p<0.001)). (H) *Tg(fli1:EGFP)* embryos at 82 hpf show increased filopodia formation in *amotL2a/b* mutants. At 82 hpf, mutants display excessive peripheral outgrowths compared with controls. (I) Quantification of filopodia length (violin plots displaying data distribution with median and quartiles, n = 10 embryos, 8 cells per embryo, N = 3 independent experiments, Mann–Whitney U test, **p<0.01). (J) Quantification of endothelial outgrowth number (violin plots displaying data distribution with median and quartiles, n = 10 embryos, N = 3 independent experiments, Mann–Whitney U test, ***p<0.001).

To further investigate how AmotL2 affects tissue compaction and remodelling, we quantified endothelial cell rearrangements as a measure of collective movements. Time-lapse imaging revealed significantly reduced endothelial cell shuffling in *amotL2* mutants, indicating deficient neighbour exchange and impaired cell rearrangements (Fig. 4E–F). Cell tracking confirmed decreased cell displacement relative to neighbouring cells, indicating that AmotL2 promotes coordinated endothelial intercalation within the vascular network (Fig. 4E–G). In control embryos, such active cell rearrangements accompany polarity fluctuations and were required for regression of low-flow branches (Fig. 4A-E), consistent with previous reports that pruning involves endothelial migration from regressing to neighbouring vessels (Franco et al., 2015; Korn and Augustin, 2015; Lenard et al., 2015). These data indicate that AmotL2 links polarity dynamics to cell rearrangement, thereby enabling flow-guided collective movements necessary for both pruning and tissue extension. Loss of AmotL2 disrupts this coordination, leading to tissue compaction, impaired elongation, and failure of branch regression.

At later stages, *amotL2a/b* mutants exhibited increased filopodia formation and ectopic peripheral outgrowths resembling lamellipodai extensions, indicative of disrupted cytoskeletal organisation and aberrant exploratory behaviour (Fig. 4H–I). These abnormalities further support that AmotL2 translates haemodynamic and mechanical inputs into cytoskeletal and behavioural responses that sustain coordinated remodelling. Together, these findings position AmotL2 as a central integrator of mechanical and haemodynamic signals during vascular remodelling. By coordinating junctional organisation, polarity dynamics, and collective cell rearrangements, AmotL2 enables flow-dependent pruning and plexus extension. The observed defects in cell polarity, motility, and cytoskeletal organisation in *amotL2* mutants suggest that AmotL2 operates at the interface between mechanical sensing and intracellular signalling.

### AmotL2 Regulates Flow-Dependent Yap Activation to Balance Vascular Stability and Remodelling

Previous work has shown that AmotL2 and Yap1 form a mechanosensitive complex at endothelial junctions. Under static flow conditions, AmotL2 sequesters Yap1 at the membrane, whereas shear stress promotes Yap1 nuclear translocation (Nakajima and Mochizuki, 2017; Nakajima et al., 2017). We next asked whether AmotL2 regulates vascular remodelling through Yap1-mediated responses to flow. To investigate this relationship during vascular expansion and remodelling *in vivo*, we examined Yap1 localisation within the sub-intestinal vein plexus. In control embryos, nuclear Yap1 was prominent in stable, maintained branches, whereas regressing branches showed lower overall levels of predominantly cytoplasmic Yap1 (Fig. 5A–B), reflecting distinct flow regimes and mechanotransduction states during remodelling.

**Figure 5.**
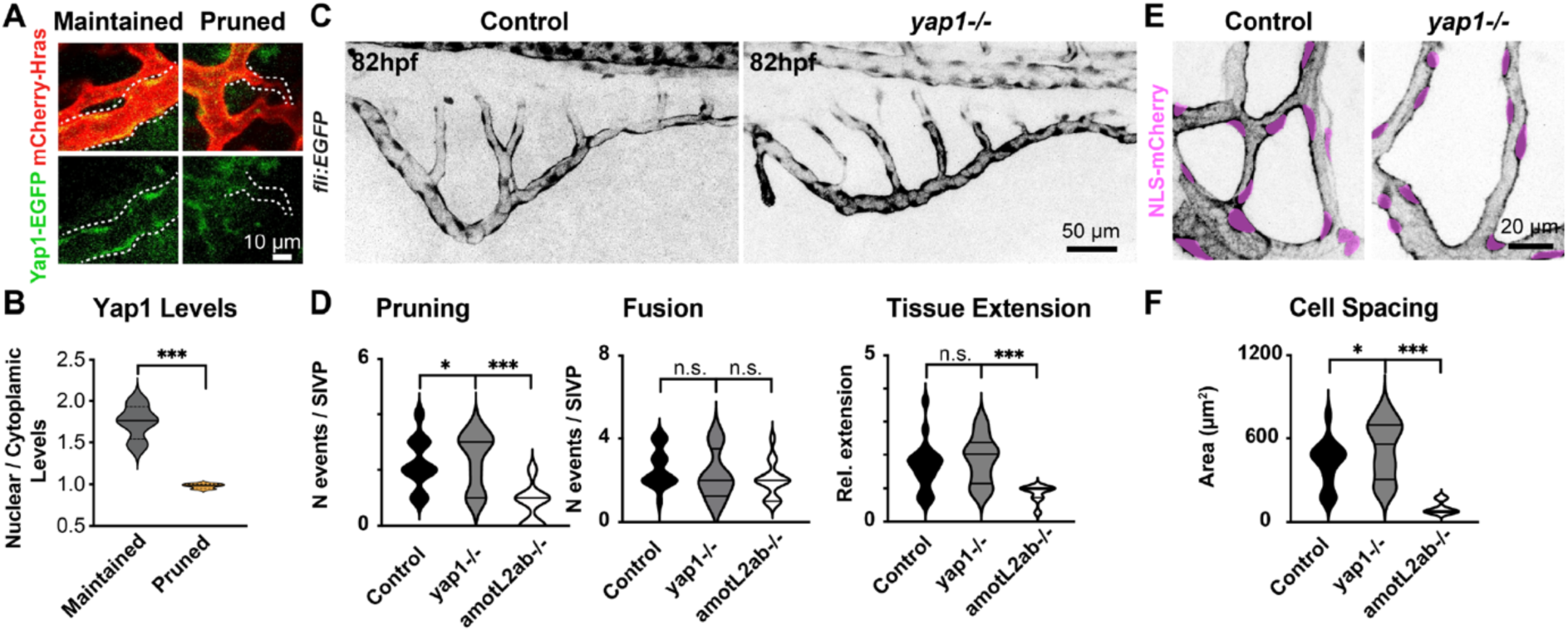
Yap1 counterbalances Amotl2 to regulate endothelial remodelling dynamics. (A) Confocal images of *Tg(fli1a:Yap1-EGFP)* embryos at 58 hpf showing Yap1 localisation within SIVP branches. Stable branches display prominent nuclear Yap1, whereas regressing branches exhibit reduced overall Yap1 levels with predominantly cytoplasmic localisation. (B) Quantification of Yap1 intensity, plotted as nuclear-to-cytoplasmic Yap1 levels, shows significantly higher nuclear Yap1 in maintained compared with pruned branches (n = 10 branches, N = 3 independent experiments, Mann–Whitney U test, **p<0.01). (C) Confocal images of *Tg(fli1a:EGFP) yap1* mutants at 82 hpf reveal enhanced vascular remodelling with increased pruning characterised by increased pruning and simplified network architecture, opposite to the *amotl2a/b* mutant phenotype. (D) Quantification of pruning events, fusion events and tissue extension in *yap 1* mutants. *yap1* mutants display slightly elevated pruning relative to controls and show a phenotype opposite to *amotl2* mutants, which exhibit reduced pruning and tissue extension (n = 10 SIVPs from 10 embryos, N = 3 independent experiments, Kruskal–Wallis test, *p < 0.05, ***p < 0.001). (D) At 70 hpf Tg(*fli1a:NLS-mCHerry*) *yap1* mutants show a modest reduction in endothelial cell density compared to controls. (E) Quantification of cell spacing in *yap1 mutants* at 70 hpf demonstrates slightly lower cell density compared with controls, and substantially lower density relative to *amotl2* mutants (n = 8 SIVPs from 8 embryos, N = 3 independent experiments, Kruskal–Wallis test, *p < 0.05, ***p < 0.001).

To assess the role of Yap1 during SIVP development, we examined *Tg(fli1:EGFP) yap1* mutants. These mutants displayed a phenotype opposite to *amotL2a/b* mutants, with enhanced pruning and simplified vascular networks, while tissue extension and fusion remained normal (Fig5. C-D). Further analysis of nuclear proximity and endothelial cell density confirmed slighly increased cell spacing in *yap1* mutants (Fig5. E-F), consistent with elevated regression activity and efficient remodelling.

Together, these observations suggest that AmotL2 and Yap1 operate in a coordinated pathway to balance endothelial stability and remodelling. AmotL2 functions as a flow-sensitive regulator, controlling Yap1 nuclear translocation and thereby linking mechanical cues to gene expression programs that maintain vessel integrity and morphology. Yap1 appears to counterbalance AmotL2-dependent stabilisation, supporting dynamic endothelial rearrangements during vascular remodelling.

## Discussion

Vascular networks must continuously balance stability with plasticity to adapt to changing mechanical environments. Here, we identify a mechanistic cascade through which haemodynamic forces shape pruning and fusion events during intestinal vascular remodelling. By integrating high-resolution live imaging, quantitative topology analysis and targeted perturbations, we show that local flow heterogeneity specifies pruning sites, that endothelial cells execute an ordered programme of polarity fluctuations and directed rearrangements to resolve unstable connections. This process is regulated by the AmotL2–Yap module which integrates shear- and tension-dependent cues to determine whether a branch is stabilised or eliminated.

### Flow as a Local Determinant of Pruning Decisions

A central finding from our work is that *local* flow differences, rather than global reductions in perfusion, dictate where pruning occurs within the SIVP. This is supported by our tricaine experiments. Despite globally lowering flow, tricaine does not enhance pruning, indicating that regression is not triggered simply by uniformly reduced shear. Previous studies in the retina, skin and brain demonstrated that low-flow segments are prone to regression (Chen et al., 2012; Franco et al., 2015), while hierarchical network models emphasise geometric determinants of shear stress (Bernabeu et al., 2014; Edgar et al., 2021). Our analysis reconciles these perspectives by showing that vascular geometry and local haemodynamics combine to create spatially patterned mechanical niches that predict pruning susceptibility. Segments with high angularity, asymmetric bifurcations or restricted lumen expansion experience disproportionately reduced *relative* flow, and these regions correlate with future regression. This interpretation is further supported by mathematical modelling, which demonstrates that local force and flow profiles adapt dynamically to the underlying network geometry (Ye and Phng, 2023). Thus, branch fate is encoded by local haemodynamic–geometric interactions, rather than by absolute perfusion level or vascular hierarchy position within the network.

### Fluctuating Polarity Orchestrates Vascular Regression

Our live imaging resolves the temporal sequence of endothelial behaviours that mediate regression *in vivo* with an unprecedented level of detail. Pruning is initiated by fluctations in endothelial polarity, followed by junctional reorganisation, redistribution of actomyosin tension and ultimately directed migration of endothelial cells out of the regressing segment. This cascade positions pruning as a dynamic, plastic process rather than a binary polarity-reversal event.

This mechanism differs from the canonical “reverse anastomosis” model derived from retina studies, where polarity was inferred from static images and interpreted as orientation against the branch toward which cells migrate (Franco et al., 2015). In our observations, polarity in regressing branches fluctuates prior to regression. In additon, under conditions of absent flow or AmotL2 deficiency, endothelial cells remain stably apolar and are often oriented at non-axial angles. Stable branches generally exhibit consistent polarity, whereas pruning branches do not. These observations indicate that polarity assessed from a single static image does not reliably reflect the underlying flow regime, and using polarity alone as a proxy for flow direction is therefore only valid in high-flow vessels, a critical consideration for studies that have inferred flow from static polarity measurements (Franco et al., 2016).

Overall, these findings demonstrate that polarity dynamics, rather than polarity direction per se, are essential for effective pruning. They also support prior studies and models (Chen et al., 2012; Korn and Augustin, 2015; Lenard et al., 2015) suggesting that regression in developing vascular networks is primarily driven by endothelial cell migration rather than apoptosis, while providing a detailed temporal framework for migration-based pruning events.

### AmotL2 Coordinates Junctional Mechanics and Endothelial Remodelling

Our data identify AmotL2 as a central regulator of endothelial mechanical competence. Consistent with its known role in linking junctions to radial actin fibres (Huang et al., 2007; Hultin et al., 2014; Hultin et al., 2017), AmotL2-deficient ECs display compromised junctional organisation and impaired capacity to remodel under differential flow. These defects manifest as disrupted cell rearrangements, reduced polarity robustness and diminished branch pruning, demonstrating that AmotL2 functions not merely as a mechanosensor but a junctional integrator of mechanical inputs, coordinating shear- and contractility-dependent inputs to productive cell movement.

Loss of AmotL2 also results in persistent tissue compaction, indicating that it maintains tension homeostasis by coupling external mechanical inputs to intrinsic contractility. By orchestarting junctional architecture, cell packing and collective rearrangements, AmotL2 ensures that endothelial cells remodel appropriately in response to spatially patterned forces. Together, these findings position AmotL2 as a central interpreter of junctional mechanics during vascular remodelling.

### Yap1 as a Counterbalancing Force That Maintains Network Stability

Our analyses of Yap1 reveal a counterbalancing function to AmotL2. While AmotL2 tunes mechanical responsiveness at junctions, Yap1 maintains normal levels of endothelial density, migration capacity and stability thresholds. Consistent with previous work (Nakajima et al., 2017), Yap1 localises to the nucleus in high-flow, stable segments and becomes cytoplasmic in low-flow, regressing branches. Loss of Yap1 results in increased, simplified networks and reduced endothelial density. These observations support a counterbalancing relationship between AmotL2 and Yap1. Taken together, our findings support a unified model in which flow magnitude and vascular geometry generate spatially resolved mechanical landscapes, interpreted through the coordinated activities of AmotL2 and Yap1. AmotL2 provides the structural capacity for endothelial cells to polarise, redistribute tension and migrate, whereas Yap1 adjusts transcriptional stability thresholds that prevent excessive or inappropriate regression. This interplay provides a mechanistic basis for how vascular networks preserve structural coherence under fluctuating haemodynamic loads.

### Integration of AmotL2 with Juncitonal Mechanotransducers

Our results highlight broader questions about how mechanical and transcriptional cues are integrated during vascular remodelling. AmotL2 imparts the ability to reorganise junctions and promote cell rearrangements, while Yap1 adjusts the transcriptional programmes that stabilise or destabilise branches. How these activities intersect with other mechanotransductive pathways is an open question. In particular, junctional mechanosensory complexes such as PECAM1, VEGFR2–VEGFR3 and VE-cadherin–associated structures are known to mediate flow sensing and endothelial rearrangement (Conway et al., 2013; Tzima et al., 2005). Understanding how AmotL2 interfaces with these complexes will be essential to define the hierarchy and coordination of force-sensing modules. Direct measurements of junctional forces using tension reporters or live force sensors, will be essential to quantify how how local loads change during pruning.

The precise mechanisms through which Yap1 regulates endothelial density, whether through proliferation, migration or cell size modulation, remain to be defined. Our findings on density are consistent with previous studies showing that Yap/TAZ integrates shear stress and cellular crowding to regulate endothelial quiescence and tissue organisation (Nakajima et al., 2017; Neto et al., 2018).

An additional question is whether mechanical thresholds differ between arterial and venous contexts. Arterial vessels experience higher shear stress and pulsatility (Red-Horse and Siekmann, 2019), which may bias AmotL2–Yap1 dynamics toward junctional stability, whereas venous beds or low-flow vascular plexi, such as the SIVP, may be intrinsically more permissive to regression. Defining how these distinct mechanical regimes intersect with actomyosin contractility to determine whether junctions stabilise or yield remains an important avenue for future investigation.

### An Integrated Model for Flow-Guided Pruning Versus Stability

Our integrated model takes into account several key features of vascular remodelling: (a) the directionality of endothelial cell movements, (b) the reproducibility of regression hotspots and (c) the preservation of high-flow conduits essential for network function.

Flow-remodelling imbalances are a hallmark of multiple vascular pathologies, including arteriovenous malformations, tumour vessel instability and impaired angiogenesis (Chiu and Chien, 2011; Furtado and Eichmann, 2024; Jin et al., 2017; Korn and Augustin, 2015; Park et al., 2021). Our observations on pruning and stabilisation dynamics may therefore have broader implications for vascular remodelling during tissue repair, vascular co-option in cancer (Kuczynski et al., 2019) and the maladaptive pruning or destabilisation seen in the cerebral vasculature during hypertension (Apaydin et al., 2022). Identifying the AmotL2–Yap1 module as a mechanosensitive remodelling node provides potential therapeutic entry points to selectively tune pathological pruning or stabilisation.

## Conclusion

Flow-mediated network refinement relies on the integration of haemodynamic cues, vascular geometry and mechanotransductive logic. By defining how AmotL2 and Yap1 cooperate to calibrate endothelial mechanics and transcriptional stability, we uncover a mechanosensitive balancing module that ensures that pruning occurs only when and where appropriate. This framework offers a unifying perspective on how mechanical forces sculpt vascular architecture during development and repair.

## Methods

### Zebrafish lines and maintenance

Zebrafish (*Danio rerio*) were maintained under standard conditions at 28.5°C (Aleström et al., 2020) in accordance with federal guidance and approved by the Kantonales Veterinäramt of Kanton Basel-Stadt. Embryos were staged according to hours post fertilisation (hpf). The following transgenic lines were used: *Tg(fli1a:EGFP)^ƴ1^*(endothelial marker) (Lawson and Weinstein, 2002), *Tg(gata1a:dsRed)^sd2^* (erythrocyte marker) (Traver et al., 2003), *Tg(fli1a:pecam1a-EGFP)^ncv27^*(junctional marker) (Ando et al., 2016), *Tg(fli1a:NLS-mCherry)^ubs10^* (Heckel et al., 2015)*, Tg(kdrl:NLS-EGFP) ^ubs1^*(Blum et al., 2008) (nucelar markers), *Tg(fli1a:B4GALT1-mCherry*)*^bns9^* (golgi marker) (Kwon et al., 2016), *Tg*(*fli1*:*EGFP-YAP*)*^ncv35^*(yap1 marker) (Nakajima et al., 2017).

### amotl2a, amotl2b and yap1 mutant lines

The *amotl2b* mutant line was generated for this study using CRISPR/Cas9-mediated mutagenesis. Guide RNAs (IDT) were designed to target exon 2 of *amotl2b*. Sanger sequencing identified the *amotl2a^ubs68^*allele harboring a six-nucleotide deletion in exon 2. Previously described mutant lines were also used in this study, including *amotl2a^fu4^*^5^ (Agarwala et al., 2015) *and yap1^ncv101^* (Nakajima et al., 2017). Homozygous *amotl2a/b* double mutants were obtained by intercrossing double heterozygous adults, and offspring were genotyped by allele-specific PCR on fin or tail biopsies.

### Immunofluorescence

Immunofluorescence staining of embryos was performed as previously described (Kotini et al., 2022) with few modifications. Embryos were fixed at 58 hpf in 2% paraformaldehyde overnight at 4°C, washed in PBS with 0.1% Tween-20 (PBST), and permeabilized with 0.5% Triton X-100 for 30 minutes. Blocking was performed in 5% bovine serum albumin (BSA) in PBST for 1 hour at room temperature. Primary antibody incubation was carried out overnight at 4°C with mouse anti–ZO-1 (1:200, Invitrogen). After washing, embryos were incubated with anti-mouse Alexa Fluor–conjugated secondary antibodies (1:500, Invitrogen) for 2 hours at room temperature. Embryos were mounted in 0.7 % low-melting-point agarose and imaged using confocal microscopy.

### Pharmacological treatments

For acute haemodynamic and cytoskeletal perturbations, embryos were treated 1 hour prior to imaging. For long-term perturbations, reatments were initiated either before the onset or at the peak of remodelling (42–58 hpf) and maintained until the end of imaging. The following reagents were used: tricaine (0.32%, Sigma) to reduce blood flow (Lenard et al., 2013), norepinephrine (60 µM, Sigma) to increase cardiac output (Chen et al., 2012; De Luca et al., 2014) and Y-27632 (100 µM, ROCK inhibitor, Sigma) to reduce actomyosin contractility (Angulo-Urarte et al., 2018). For combined treatments, tricaine and Y-27632 were co-applied at the same concentrations. Embryos were maintained in E3 medium containing the respective compound throughout imaging.

### Imaging and image analysis

Live imaging was performed using a Zeiss LSM880 confocal microscope equipped with Airyscan or an Olympus SpinSR spinning disk confocal system. Whole SIVPs were imaged using a 25X oil-immersion objective (NA 0.8), while focused regions were acquired using a 40X silicon oil-immersion objective (NA 1.2) or a 30X oil-immersion objective (NA 1.05, SpinSR). Fixed or live embryos were selected based on fluorescence signal, anaesthetised in E3 medium containing tricaine (0.08%, pH 7.0, Sigma) and mounted using 0.7% low-melting-point agarose (Sigma) on glass-bottom dishes (MatTek). Imaging was performed under temperature-controlled conditions (28.5°C).

Flow directionality was determined by tracking *gata1a:dsRed* positive erythrocytes. Microscopy datasets were stored and managed on OMERO platforms and processed using Fiji/ImageJ and Imaris (Bitplane) software. OMERO was also used for image organisation and figure preparation.

### Quantification and statistical analysis

#### Vessel geometry and remodelling analysis

Vessel geometries were classified as L-type (pruning-associated) or O-type (fusion-associated) based on 3D topology of connected vessel loops. The proportion of L- and O-type segments undergoing pruning or fusion was quantified manually from time-lapse datasets. Morphogenetic events (pruning, fusion, sprouting, anastomosis) were quantified as events per SIVP, while tissue extension was expressed as the relative increase in total SIVP area from the initial timepoint (within a 12h period).

#### Diameter analysis

The pruning diameter threshold was defined as the mean vessel diameter immediately preceding regression in control-pruned branches (∼4 µm). The per-embryo pruning proportion was calculated as the fraction of below-threshold branches that subsequently regressed.

#### Cell density analysis

Endothelial nuclei in *Tg(fli1a:NLS-mCherry)* embryos were segmented in Imaris to extract nuclear centroids. Local cell density was then estimated by computing Delaunay triangulation from these centroid coordinates using existing Fiji scripts. For each nucleus, the areas of the Delaunay triangles formed with its nearest neighbours were measured. These triangular areas provide a robust, geometry-based estimate of local cell spacing, where smaller areas indicate increased cell density and tissue compaction.

#### Polarity and cell shuffling analyses

Endothelial polarity was quantified in *Tg(kdrl:NLS-EGFP; fli1a:B4GALT1-mCherry)* embryos. Golgi–nuclear angles relative to flow direction were determined using nuclear centroids segmented in Fiji, and polarity variance was computed as the circular standard deviation of angular displacement over time. Cell shuffling was quantified as the number of neighbour exchanges per branch over ∼5 h of imaging. Nuclei were segmented and tracked using TrackMate (Fiji). A neighbour exchange was defined as a cell gaining and losing at least one neighbour between consecutive timepoints. The per-branch shuffling rate was calculated as the mean number of neighbour exchanges per cell within the 5-hour timeframe.

#### Filopodia and outgrowth analyses

Filopodia length and peripheral outgrowth number were measured in *Tg(fli1a:EGFP)* embryos. Images were background-subtracted, small z-stacks were maximum-projected, and vascular peripheries were skeletonised in Imaris. Filopodia were identified using line ROIs in Fiji, and the mean filopodia length per cell was calculated for each embryo. Peripheral outgrowths were quantified manually at 80–82 hpf per embryo.

#### Sampling and experimental design

All imaging and quantification were performed using unbiased random sampling of embryos from each clutch. Pharmacological treatments were randomly allocated across embryos within each clutch, and imaging and analysis were performed without pre-selection based on phenotype. For genetic experiments, mutant lines were maintained as heterozygous crosses and analyses were conducted blind to genotype, followed by post hoc genotyping of individual embryos to confirm classification.

#### Statistical analysis

For all analyses, each data point represents one embryo (biological replicate) corresponding to a single SIVP. Sample sizes, exact statistical tests, and p-values are reported in figure legends. Analyses were performed using Fiji, R, and GraphPad Prism10. All quantifications were performed from ≥3 independent experiments, with biological replicates as indicated. Statistical analyses used non-parametric tests (Mann–Whitney U or Kruskal–Wallis with Dunn’s multiple comparisons) unless otherwise specified. Data are presented as mean ± SEM or median with interquartile range, and significance was set at p < 0.05.

## Acknowledgments

We thank Kumuthini Kulendra and Andre Rodriguez for fish care and the Imaging Core Facility of the Biozentrum (University of Basel) for microscopy support. We also thank Naoki Mochizuki for essential zebrafish lines and critical reading of the manuscript. This work has been supported by the Kantons Basel-Stadt and Basel-Land and by grants from the Swiss National Science Foundation (310030_200701 and 310030B_176400) to M.A. M.P.K was also financially supported by the Junior Research Fund from the University of Basel.

## Author contributions

M.P.K. and M.A. conceived the idea and directed the work. M.A. provided supervision and financial support. M.P.K. designed and performed experiments. L.M. and E.S. performed experiments. M.P.K. and M.A. analysed the data. H.N. provided essential zebrafish lines and reagents. M.P.K. wrote the manuscript. All authors reviewed the manuscript.

